# Involving patients and publics in medical and health care research studies: an exploratory survey on participant recruiting and representativeness from the perspective of study authors

**DOI:** 10.1101/410480

**Authors:** Jonas Lander, Holger Langhof, Marie-Luise Dierks

**Author notes:** Corresponding author: E-Mail (JL).

## Abstract

Research on patient and public involvement so far concentrates on defining involvement, describing involvement methods, and analyzing involvement practice in various individual research disciplines. There is little empirical data on the process of and aims for selecting participants, and to what extend lay people involved in research can and should be representative of the population at large. To explore practices and perceptions on these issues and on future PPI conduct more generally, we sent an electronic survey to authors who published involvement activities as part of their studies in medical and social science journals. We identified such authors with a systematic search of five databases and applied descriptive statistics for analysis. Of those who returned the survey (n=127 of 315; 40%), most had previously conducted involvement activities (73%). 45% reported more than one type of involvement, e.g. consultation and deliberation and participation (14%) and to have recruited more than one type of participant for their PPI activity (56%), e.g. ‘lay publics’ and ‘expert publics’ (33% of 71). Representativeness was seen by most respondents as a crucial objective when recruiting participants, while less than half found the recruitment of suitable participants very easy (9%) or rather easy (34%). Many respondents found it generally difficult (52%) or very difficult (17%) to achieve good representativeness. They identified significant respective challenges and desired more guidance on various aspects of planning and conducting PPI (56%). 55% thought that the concept of “involvement” should be changed or improved. We conclude that participant recruitment and representativeness are controversial in current PPI practice given the manifold challenges mentioned by the survey respondents. Our findings may inform further research particularly regarding– the potentially large number of – unpublished PPI activities.

## Introduction

Recent research on patient and public involvement (PPI) has identified a range of reasons to involve patients and publics in medical research and health policy. These include the need to align research priorities with societal preferences [1–3], the reorganization of existing health care services [4], and the need to assess the impact and value of health technologies and health services [5]. PPI may also increase transparency, legitimacy and accountability of scientists and policy-makers vis-à-vis society [6–8]. Others highlight its importance simply because so many decisions about health care are financed by tax payers, who should therefore have a stake in relevant decision-making processes [9].

Various (inter-)national organizations such as INVOLVE UK, NICE Patient and Public Involvement Programme (PIP) and the International Association for Public Participation (IAP2) have established PPI advisory and interest groups. These groups serve different objectives and tasks, mainly advancing PPI frameworks, strengthening PPI in guideline development, and making PPI a prerequisite for instance for project funding. Another common issue is the actual definition of PPI. While full consensus is yet missing, there is a tendency to distinguish several stages of active and passive involvement, i.e. a) informing/educating (least active), b) consulting for opinions and preferences, or c) inviting lay people to participate in planning and conducting research (most active) [1,10,11].

Besides such more general, conceptual issues, there are particular challenges for realizing involvement and deciding whom to involve. An initial challenge concerns the distinction between ‘patients’ and ‘public’ [12,13]. For example, patients may want to influence the setting of health research priorities to benefit their individual health. ‘Publics’ may be more concerned about the amount of money invested in medical research generally. It is further argued that ‘public’ often lacks clarity, and that too many terms such as ‘citizens’, ‘service users’, ‘community members’, ‘lay people’, ‘carers’, and ‘tax-payers’ are used without explication [8,14]. While ideally there should be clear distinctions, this may often be difficult in practice, and patients may be defined simply as a “subgroup of the wider role of citizen or member of the public” [13].

Related to these distinctions is the challenge of determining if and to what extend those involved in research should be representative of those who are not involved. On the one hand, representativeness is needed to avoid the systematic exclusion of some social groups [8]. Also, because the usually small number of participants influence decisions made on behalf of many others, they must be carefully selected [15]. In cases where representativeness is necessary, researchers and other organizers need to ask whether it is more important to recruit a sample that accurately reflects socio-demographic characteristics (i.e. statistical representativeness in terms of age, gender, etc.) or one that reflects the broadest possible range of existing views (qualitative representativeness) and discourses on a given subject (discursive representativeness), or whether certain representatives should be elected to act on behalf of others (elective representativeness).

On the other hand, depending on the precise aim and subject, a PPI activity may require specific participants that cannot be regarded representative, for example patients with advanced disease-specific knowledge. Also, as long as PPI is a ‘side feature’ rather than an integral part of research, the organizational and financial resources of research teams may not allow for highly sophisticated recruiting processes that ensure good representativeness. Hence, “true representation” may be difficult to achieve in practice [16–18] and may not be desirable in each case.

Various previous studies assess and debate PPI terminology and the challenges it entails [2,13,14], different types of representativeness and how they are used [17,19], and current PPI practices more broadly [20–23], using for instance debates, framework developments, and literature reviews. Few studies take a more practice-oriented approach by conducting interviews with professionals on PPI in general [24,25], and with small sample sizes [17,22]. The GRIPP2 checklist for reporting PPI in research recommends to describe and transparently define the individual steps taken during PPI, such as the definition of PPI for that study and the methods used [26].

To our knowledge, this is the first study aiming to explore practices and preferences regarding participant recruitment, handling of representativeness, and PPI conduct from the perspective of authors of published PPI activities. It also tries to shed light thereby on the debate about whether PPI is a practical concept or merely confusing, and to what degree representativeness affects meaningful involvement.

## Materials and methods

### Data search

The data were gained via an online survey distributed to corresponding authors who published their PPI activities in the field of medical and health care research. To gain the necessary contact information, we first conducted a systematic search for studies in relevant medical and sociological databases (Fig 1). For this, MeSH terms of a) PPI and b) medical research, healthcare research and health policy were combined into search queries which we adapted to the respective database where necessary (Table 1).

**Table 1.**
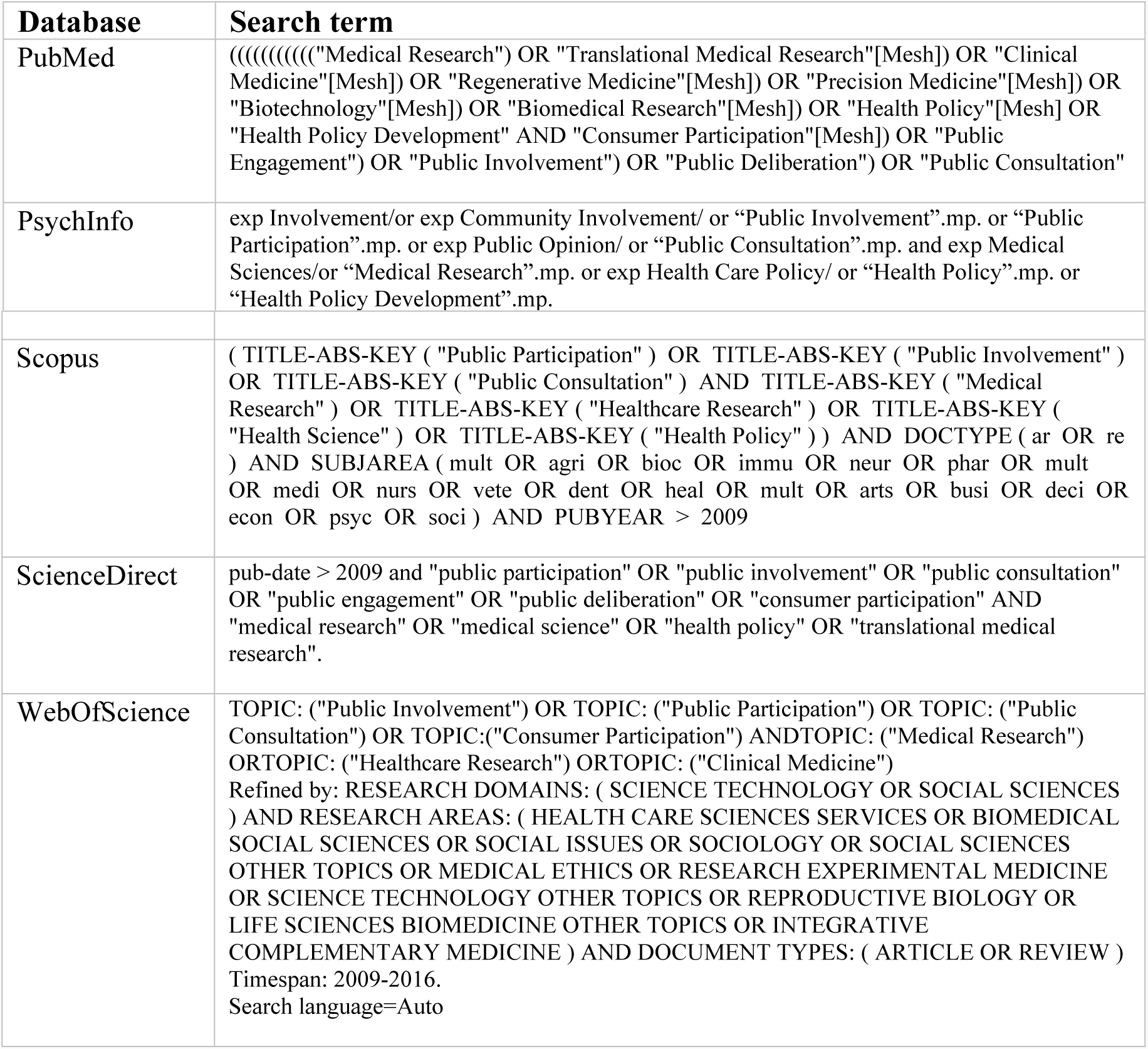
Search terms

**Fig 1.**
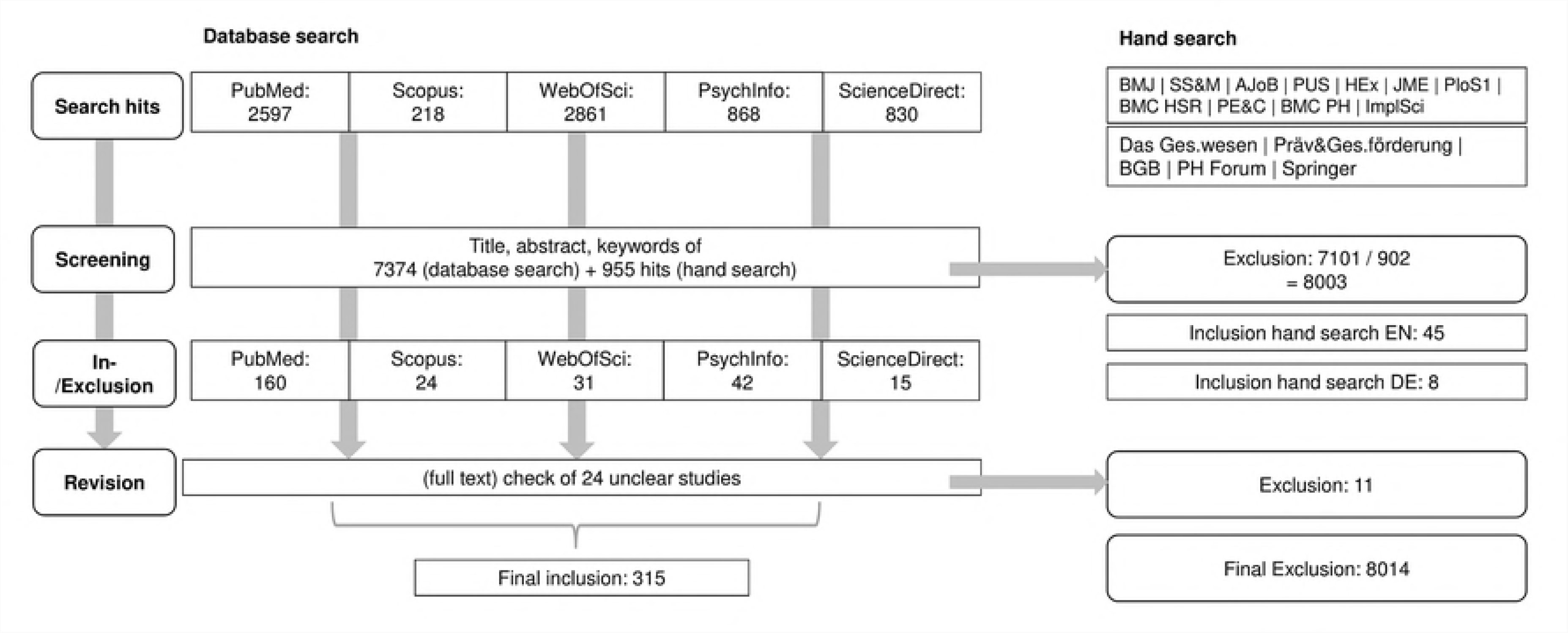
Database search

We included publications if they a) directly reported on PPI as part of a research project, b) in medical or health care research, c) between 2009 and 2016, and d) in English or German. Regarding a), we framed PPI as spectrum of activities with varying degrees of active and passive involvement (see introduction). This was done since a) different researchers and other PPI actors have different understandings of what involvement is and hence also use different terminology, e.g. involvement, engagement, participation, and deliberation, and b) the reporting on PPI is neither mandatory nor standardized yet, making it difficult to focus on a specific type of involvement alone.

A sample of 800 search hits was screened separately by each of two researchers (HL, JL) to achieve coherence regarding the in- and exclusion of studies. The screening results were compared, and minor differences were resolved by discussion. We also discussed studies that did not show clearly how the input gathered could be used to develop or adapt research or decision-making processes and excluded some. For instance, one study reported experiences of women with cancer diagnosis, but was unclear about the translation of this input into practice. We also hand-searched those journals that yielded at least 3 PPI studies in the database search (n=11).

### Data-collection

From the included studies, we collected the necessary information about the corresponding authors, including name, position, and email address. This information was updated in many cases as authors had moved institution or changed their contact details.

The survey included questions with single and multiple answers on a) authors’ previous experiences with PPI, b) specific questions on recruitment and underlying aims regarding participant characteristics and representativeness, and c) more general preferences for future PPI (S1 Table).

Authors were pre-notified of the research project via email one week before the actual survey. During an eight-week period of collection survey responses (March – April 2017), two follow-up reminders were sent to elicit further answers. Ethical approval was gained from the Ethics Commission of Hannover Medical School (Reference number 3465-2017).

### Data analysis

The data were collected via SurveyMonkey, and exported to SPSS Statistics for analysis of frequencies, correlations between answers to different questions – for instance recruiting aim and recruiting methods – and assessment of multiple answers (using multiple answer sets and cross tables). Since “involvement”, “participation” and “deliberation” may be more ‘active’, ‘direct’ or ‘ideal’ types of PPI than public “consultation”, we analyzed the most relevant issues separately for these types of PPI and mention differences where applicable. An a priori quality assessment of selected studies was not applicable since the focus was on authors’ practices, not on the reporting.

## Results

### General characteristics of studies

Of 8,329 search hits, 315 were eventually selected based on the criteria stated in 2a), and we attempted to contact their corresponding authors. The final response rate was 40% (n=127). Since the survey was anonymous unless authors voluntarily provided the study title, details regarding place, year of conduct and subject of the involvement activity could only be obtained for 65 studies (Table 2).

**Table 2.** General information about included studies

|  | Subject <sup>a</sup> | n | PPI aims | n | PPI methods | n | Recruited participants | n | Number of participants | n | Duration of participation (days) | n |
| --- | --- | --- | --- | --- | --- | --- | --- | --- | --- | --- | --- | --- |
| a) | Genetics, genomics | 10 | Consultation (only) | 40 | Questionnaire (only) | 16 | Lay publics (only) | 26 | Up to 10 | 10 | Up to 1 | 78 |
| b) | Biobanking, biobank research | 6 | Deliberation (only) | 7 | Interview (only) | 6 | Expert publics (only) | 4 | 11-30 | 25 | 1-2 | 13 |
| c) | Priority setting in research and/or healthcare | 3 | Participation (only) | 8 | Focus group (only) | 8 | Lay patients (only) | 13 | 31-50 | 15 | 2-5 | 30 |
|  |  |  |  |  |  |  | Expert patients (only) | 6 |  |  |  |  |
| d) | Cancer research, cancer treatment | 3 | Consultation and deliberation | 8 | Assessment (only) | 0 | Family members or representatives or medical staff (only) | 1 | 51-100 | 14 | > 5 | 0 |
| e) | Research, clinical trials (not subject-specific) | 4 | Consultation and participation | 10 | Citizen jury (only) | 1 | Others (only) | 6 | 101-200 | 10 |  |  |
| f) | Decision-making and change | 3 | Other combination with 2 aims | 7 | Questionnaire and interview | 10 | Lay and expert publics | 4 | 201-500 | 18 |  |  |
| g) | End-of-life care | 2 | Consultation and deliberation and participation | 8 | Questionnaire and focus group | 9 | Lay and expert patients | 6 | More than 500 | 30 |  |  |
| h) | Quality, quality indicators | 2 | Information and consultation and deliberation | 8 | Other combinations with 2 methods | 21 | Other combinations with 2 types of participants | 9 |  |  |  |  |
| i) | Research participation | 2 | Information and consultation and deliberation and participation | 8 | Information and questionnaire and interview and focus group | 4 | Lay publics and expert publics and lay patients and expert patients and family members and representatives and medical staff | 5 |  |  |  |  |
| j) | Other subjects <sup>b</sup> | 29 | Other combinations with 3 or more aims | 14 | Other combinations including 3 or more methods | 47 | Other combinations including 3 or more types of participants |  | 40 |  |  |  |

<sup>a</sup> only those studies for which the authors voluntarily mentioned the study title (n=65)
<sup>b</sup> Autism research, Long-term conditions, Drug licensing, Pharmacy alcohol screening, Depression, Consent for emergency research, Rheumatology, Surgical research, Grief, Ageing care, Dietetic consultations, Drug advertisement, Vaccine safety, Intermediate pulmonary nodule, Cerebral palsy, Diabetes, Organ donation, Nanotech, Dementia, Sensitive health topics, Health tech design, Mental health, Dengue control, Embryo status, Degenerative ataxia treatment, Biomedical research, General practice, Physician-pharmacy interaction, Health insurance

Of the studies with known titles, 10 reported genetics/genomics as the subject. Biobank research (n=6) and research not associated with a specific discipline (n=4) were also reported repeatedly. Three subjects were reported three and two times respectively, including priority setting (n=3), decision-making and change (n=3), and end-of-life care (n=2). The rest (n=24) were reported only once (Table 2).

The majority of authors had conducted PPI before (73%); most of them (46%) had been involved in between two and five PPI activities. 21 authors said they had already conducted six to ten or even more activities.

Across all 127 studies, consulting participants for preferences, opinions, etc. was reported most often as the sole aim (31%). Deliberation and participation were reported seven and eight times respectively as the sole aim. All other studies reported a combination of aims, such as consultation *and* participation (8%), or information *and* consultation *and* deliberation *and* participation (6%). While deliberation and participation were rarely reported alone, they were often reported alongside other aims (deliberation 39%; participation 33%).

The most often reported single methods used to achieve these aims were questionnaires (13%), interviews (5%), and focus groups (6%). Information provision, assessments and citizen juries were only used in combination with other methods, e.g. focus groups. The most frequent combinations were questionnaires and interviews (8%), questionnaires and focus groups (7%), and information *and* questionnaires *and* interviews *and* focus groups (6%).

Among the various involvement aims, the distribution of methods was consistent, i.e. most often questionnaires/interviews, then focus groups, information, and assessments/juries, with “other” methods in last place (Fig 2a). The use of more active involvement methods such as assessments and juries as well as “other” methods (e.g. “deliberative polling”, “reviewing of research material”, “participatory workshops”) was higher for deliberation/participation studies (assessments = 8%; juries = 21%; other = 25%) than for consultation studies (5%; 5%;5%).

**Fig 2.**
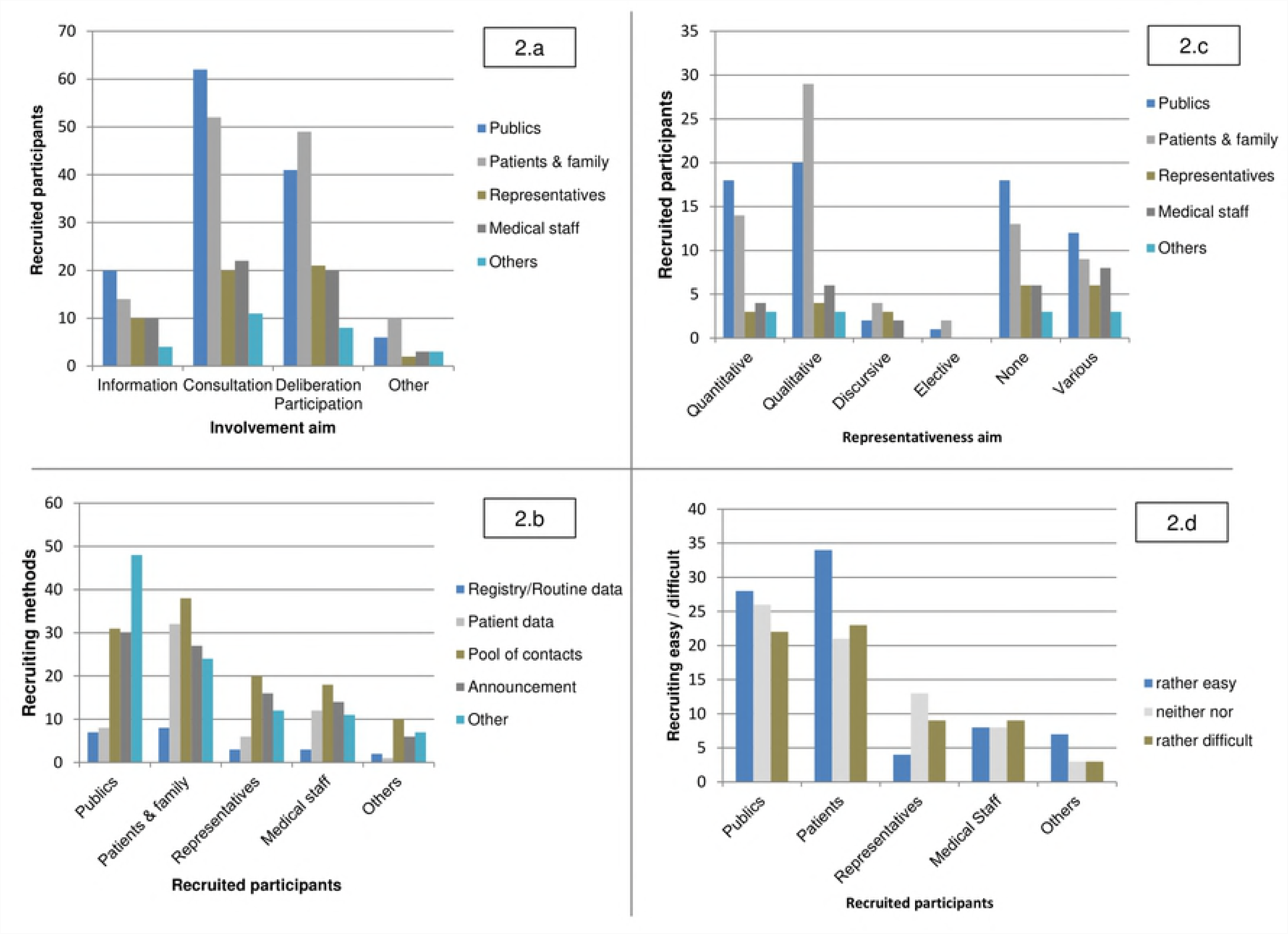
Summary of results

### Aims for recruiting participants and handling representativeness

50 studies recruited only one type of participant, most often lay publics (20%) or lay patients (10%). Studies that recruited only one of expert publics and patients, family members and representatives were much rarer. 71 reported recruiting a combination of two or more. These included lay and expert publics or patients, and lay and expert patients with family members and representatives. 5 studies reported a combination of 6 different types of participants, such as lay and expert publics, lay and expert patients, family members, and representatives. All other combinations were reported once each (Table 2).

Recruitment methods were similar for the various participants: most often from a pool of contacts, then announcements, patient data, “other” methods, and least often registry/routine data (Fig 2b). Only for recruiting lay and expert publics were “other” methods used almost twice as often as when recruiting patients, representatives, etc.; for instance, “speaking at community events”, “call for interest” in community groups, and social media.

Many authors rated representativeness of the participants vis-à-vis the general population very (33%) or rather important (38%). Studies that aimed solely at consultation (34%) or deliberation/participation (18%) showed no particular difference regarding the importance of representativeness compared to the full sample. The two subgroups (consultation and participation/deliberation) accorded representativeness equal importance. 17% of all authors, however, indicated that representativeness was rather not or not at all an important sample characteristic.

When recruiting participants, most authors either aimed for quantitative (25%) or qualitative representativeness (32%). Within the subgroup of deliberation/participation studies, slightly more studies aimed at qualitative (35%) and quantitative representation (30%) compared to the overall sample (32%; 25%). Also, deliberation/participation studies less often reported that they had no representativeness aim (13%; overall sample 20%). Discursive and elective representativeness were reported only rarely (4%; 2%) and 19% of all respondents did not indicate any aim.

Among the various representativeness aims, the distribution of (types of) recruited participants was similar: most often lay/expert publics, then lay/expert patients, medical staff, representatives, and last “others” (Fig 2c). However, lay/expert publics were reported most often for quantitative representation, and lay/expert patients for qualitative representation.

### Outcomes of the recruitment process and levels of representativeness achieved

55 respondents mentioned that recruiting the intended group was very easy (9%) or rather easy (34%). 36 found it to be neither easy nor difficult (29%), rather (28%) or very difficult (2%). The group of deliberation/participation studies included more rather/very difficult and neither–nor ratings (67%) than the overall sample (56%). In contrast, consultation studies more often rated the recruiting very/rather easy (58%) than did deliberation/participation studies (33%) and the overall sample (44%).

Recruitment was never mostly found to be easy: there were almost as many “rather/very difficult” ratings as “rather/very easy” ratings for every type of participant (Fig 2d). Only for recruiting lay/expert patients and their family members, who were most often recruited via a contact pool and patient data, did recruitment seem somewhat easier.

Apart from those respondents who reported recruitment difficulties, most indicated success in recruiting the intended participants eventually (86%). Still, 33% of respondents who answered the question “If you did not manage to recruit the intended group, did this influence the generalizability of study results?” said that the study results were not fully generalizable. Another 38% did not know whether there was any influence.

Besides the conclusions respondents drew from their respective studies, we also asked for their more general perspective on recruitment and representation. Here, 71% said that representation is generally rather (40%) or very important (31%). Only 5% found it rather or very unimportant; all others were undecided.

In contrast, actually achieving representativeness was perceived to be harder. On the question “Do you think that representation is generally possible/achievable?”, 69% viewed this as rather (52%) or very difficult (17%). Even among those who saw representation as (very) important, a considerable part indicated that it is generally rather not (n=40) or not possible to achieve (n=8).

Regarding the relevance of (future) guidance for conducting PPI, 31% think that more guidance is needed on overall planning and conduct. Guidance for recruitment was desired by 28% of respondents, who often looked to research organizations for guidance. As for the difference between consultation and deliberation/participation studies, authors of the latter desired overall guidance more often (36%) than authors of consultation studies (21%).

Further, more than half of all authors (55%) indicated that the concept of PPI as such requires modification. The authors of deliberation/participation studies agreed more often (59%) than the overall sample (55%).

Finally, 37% felt that the terms “patients” and “publics” need to be better defined or differentiated. Within the group of deliberation/participation studies, authors were happier with the terminology: only 23% would welcome changes.

### Open answers on the relevance and challenges of recruiting participants and achieving representativeness

45 of 127 respondents answered the open-ended question “Based on your experiences, is there something regarding the selection of participants or the role and relevance of representativeness of participants that you want to add here?” The analysis allowed for a topic-wise categorization of answers (S2 Table).

The greatest number of comments (33%) was made regarding challenges and limitations of participant selection and the feasibility of representativeness. Authors mentioned for instance how hard it is to “control generalizability [of study results] completely”, as participant selection can always include some self-selection. Regarding representativeness, it was stated for instance that “everyone [has] a different view of what representative participants would be”, “the need for representation varies based on [different] studies”, and “I don’t think that true representativeness is possible” (Box 1).

##### Box 1. Extracts from free-text answers Representativeness

“The importance of representativeness of participants strongly depends on you project/research objectives”

“It is essential to high quality research. There is much guidance on PPI already available, however, there is no harm in receiving more to ensure we do it better”

“The notion of representativeness in PPI is a load of nonsense and entirely misses the point. Talking about representation deflects from the real value of PPI. No one questions how representative I the researcher am when I conduct a project. Why do we need to question how representative patients are? It’s time to move away from this and get on with involving patients in our work.”

##### Recruitment

“I think it is important that any patient and public involvement be clearly identified in regard to the research questions and aims. Specifically, why do you want to speak with or engage with this group and how will their contributions help with the research.”

“It is essential that the differences between patients and publics be understood. As well, we need a third category - “community” which identifies a different public constituency with collective interests that also warrant engagement and representation.”

##### Limitations and challenges

It’s not good idea to use the same people for PPI to represent a group on different issues. The PPI group should have expertise in the area or topic, and the same people could not be in PPI.

“‘Representativeness’ should be approached with extreme caution as it is rare for people to agree on what participants should be representing: demographics, perspectives, the ‘everyman’, the experiential expert etc., and thus it can be manipulated.”

Regarding participant selection, the respondents mentioned a range of issues that demand careful consideration (see section e) in Supplementary file 2) (22%). For instance, values, orientations, and topic-specific opinions may be as important as socio-demographic characteristics when selecting participants. More generally, much more effort and “hard thinking” is needed to select participants appropriately than to simply replicate the wider socio-demographic range. It was also emphasized that it is crucial to select the sample based on the specific PPI objective, and to differentiate participants instead of mixing them up.

As for the relevance of representativeness, some authors opined that it is indeed an important prerequisite for successful PPI (see section a)). However, the majority indicated that representativeness is rather not important/relevant, or challenging in practice, not least given the difficulties of finding a common definition and implementing it. Three respondents argued that focusing on representativeness may distract researchers from other, more important issues in PPI, and that the wording/notion may easily be misleading, e.g. it “deflects from the real value of PPI”. Another respondent suggested that representativeness may simply not be desirable, for instance when being interested in “a range of perspectives” and “views” instead.

## Discussion

### Relation of main findings to previous research

Besides those previous studies that discuss recruitment and representativeness from a theoretical perspective or address them very generally in reviews, few have dealt with this subject in detail and/or using a similar methodology. [24] and [25] investigated industry and researchers’ attitudes towards PPI using interviews (n=15; n=21), though without any particular focus on recruitment and representativeness. A recent study by South et al. [27] broadly describes the kind of “representatives” involved in a small sample of UK-based case studies, though without further (quantitative) analysis of the details of recruiting or representativeness.

Others assessed how representativeness is understood in different contexts using qualitative interviews [17], and current challenges from the perspective of research network members [22]. While the latter did not address representativeness per se, they even speak of a “crisis of representation”. Further, the RAPPORT study [28] assessed current PPI in different research disciplines from the perspective of UK-based principal investigators, though without any specific focus on the handling of recruiting and representativeness.

While these studies clearly frame recruitment and representativeness as central issues, we aimed to explore how these aspects are handled in practice by one of the principal groups concerned with PPI, i.e. study authors. It may be easier to deal with these issues once actual practices are known.

The vast majority of authors had had prior experience with involving lay people in research, which may help with general aspects such as deciding on a recruiting method. Still, many authors would welcome more guidance; hence, previous experience may not automatically help researchers deal with trickier matters such as determining whether and what type of representativeness is necessary for a particular study. Researchers may also find it difficult recruiting ‘representative’ samples if they themselves belong to a specific part of society and research community, that does not per se have (easy) access to those who are more difficult to recruit, i.e. vulnerable populations.

A considerable number of studies recruited more than one target group (38% at least 3), often mixing up distinct types of participants. This may help increasing the scope of the discussion. However, one of the current challenges is to better define and differentiate whom to involve in research, and how. Also, few of those studies for which authors gave the title provided an explanation for why specific participants were included or what “public” etc. means in their particular case, other than to provide general sample information. Also, while respondents often seemed overall positive about their own study, many (Q. 19) expressed conceptual and practical difficulties with how participants differ from each other, and what representativeness means in different contexts.

Because of this difficulty and since PPI activities are not solely determined by participant representativeness, social science research argues that other concepts such as inclusiveness may be employed instead. Aspects such as individuals’ abilities and resources to influence decision-making, the organizational context that enables or hinders individuals’ participation and impact, the professionalization of participants, and the actors’ networks and changing power relations are influential on PPI, too [29–31]. This assumption is supported in different responses to our survey, stressing the boundaries of representativeness (see Supplementary file 1).

### Methodological considerations

To reach as many authors as possible, we conducted a systematic search of multiple databases. We also did not limit the search by discipline, allowing for insights into PPI practices across a range of research areas. However, we could not conduct a subgroup analysis regarding potential differences in the recruiting and handling of representativeness across different research topics as the survey was anonymous and hence, retracing all topics from the respective studies was not possible.

Further, as the reporting on PPI and particularly the rather and very active types of involvement is neither mandatory nor standardized [11], we applied a rather broad definition of involvement that includes more and less active types. Further research is needed to identify potential differences in recruiting aims and processes as well as on the perspectives on representativeness across the different types of involvement.

While it may generally be important to differentiate types of PPI, our objective was to assess recruitment and representativeness practices. This may be equally relevant both for more and less active types of involvement, as any study needs to select participants carefully. Nevertheless, PPI activities may take place in various academic and non-academic settings, and its methodology may hence differ, as well as the individuals and groups responsible for planning and conducting PPI. Since we contacted PPI organizers based on publications in medical and social science journals that directly reported on a PPI activity, we could not include involvement efforts not published along research projects. Hence, it may be vital to enhance our findings by analyzing a) those PPI activities published or reported in different formats and b) cases where PPI was part of a research project, but not published.

## Conclusion

The results of our survey highlight several challenges, namely deciding when and what kind of representativeness is required according to the study aim, defining and differentiating among different ‘types’ of participants and justifying why certain participants were chosen over others and, more generally, distinguishing between ‘patients’ and ‘publics’. The latter aspects became evident in particular from the survey respondents’ open answers (Q. 19). Also, while (some) guidance on planning and conducting PPI is available from research and PPI institutions, a considerable number of survey respondents mentioned the need for more guidance and a better understanding of “PPI”. Hence, available support may not have been a specific help for many of the study authors we surveyed.

In view of the practical difficulties of selecting participants mentioned by the survey respondents as well as the lack of reporting on PPI activities in general and regarding recruitment in specific (see below), those planning and conducting PPI may consider the following aspects: a) providing an explicit justification for why certain participants were selected and how the recruiting methods fit the recruiting aims, b) making the type and degree of representativeness of participants dependent on the study aim and not seeing representativeness as a prerequisite, c) defining the exact role of participants in the involvement process and how they are involved and thinking about potential implications for selecting participants, d) considering whether the experiences, skills and qualifications of the research team are sufficient to plan and conduct a PPI activity, and e) being clear about how potential recruiting and representativeness limitations impact on the process and results of the involvement activity.

Since our findings and the named challenges are only representative of the views and practices of study authors who published PPI activities as part of their research, further research seems relevant to assess how the named challenges are handled in those cases where PPI activities as part of research projects are not published and whether this differs from the findings presented here. Further assessment of PPI activities could be done for instance by contacting research groups leaders either from a specific area of research or more broadly and interviewing them regarding non-published PPI activities. This may also reveal insights about whether reporting/publishing on PPI and particular aspects such as participant recruiting is lacking so far [19] because authors simply do not consider reporting about PPI relevant, or because of actual difficulties with PPI conduct in practice.

Lastly, as various theoretical and practice-oriented contributions to PPI research are now available, future research may also focus on consensus-finding approaches to the pertinent challenges. In particular, PPI is still understood too variably in terms of what it means, and which methods are more or less suitable in different research contexts. The results of our research confirm this thesis, particularly regarding PPI authors’ desires for methodological guidance and for changes to or consensus over the concept of PPI itself.

## Acknowledgements

We thank Daniel Strech and Iris Brandes for their very valuable advice on developing the research question and structuring the manuscript, and Reuben Thomas for his support with language and proof-reading. We also thank Jahnavi Daru and Rachel Matthews for their helpful comments during the revision phase.

## Supporting Information

**S1 Table. Survey**

**S2 Table. Categorization of open answers to question 19**

